# ClonoDeconv: reference-based *α − β* paired recurrent clonotype proportion estimation from bulk-TCR sequencing

**DOI:** 10.64898/2026.10.13.758550

**Authors:** Yuze Zhou, Elie Antoun, Mariana Pereira Pinho, Jianqing Zheng, Yuxi Wu, George Adigbli, Fadi Issa, Yanchun Peng, Julian C. Knight, Tao Dong

## Abstract

T cell receptor (TCR) sequencing is crucial when studying antigen-specific T cell responses and clonal dynamics. A key objective of TCR repertoire analysis is to identify recurring clonotypes across timepoints or different disease states, providing insight into the evolution of immune response. Bulk-TCR sequencing enables high-throughput profiling of TCR *α* and *β* sequences but lacks pairing information, while single cell TCR sequencing (scTCR-seq) retains pairing information but is limited by sensitivity, throughput, and costs. We present ClonoDeconv, a computational framework to deconvolute paired TCR *αβ* clonotype frequency from bulk TCR sequencing experiments using a paired scTCR-seq reference. ClonoDeconv robustly estimated shared clonotype abundances from 5’ RACE bulk-TCR datasets using a scTCR-seq reference from the same donor. Applied to longitudinal samples, ClonoDeonv successfully captured the expansion and contraction of rare virus-specific clones across timepoints. Taken together, ClonoDeconv provides a cost-effective and scalable approach for studying the longitudinal T-cell clonal dynamics.

## Introduction

The T cell receptor (TCR) is predominantly a heterodimeric protein of *α* and *β* chains. Both TCR chains are generated in a random process involving V(D)J gene segment recombination and nucleotide insertion and deletion^1^. Gene segment diversity and junctional diversity within each chain, combined with the diversity introduced from alternative pairing between *α* and *β* chains, together result in an extremely diverse TCR repertoire in each individual^2^. The identity of the TCR determines antigen-specificity with the CDR3 loops of both chains together forming the binding interface with the peptide-major histocompatibility complex (pMHC)^3,4^, making an alpha/beta paired TCR molecule the fundamental unit of antigen-recognition^5,6^. Moreover, because of the extremely low probability of generating the same TCR*αβ* sequence independently twice^7^, T cells with the same TCR clonotype can generally be regarded as derived from a common ancestor, namely, they belong to a single clone. Therefore, TCR sequence can be used as a barcode to trace the expansion and contraction of T cell clones across different biological and experimental conditions^7^.

The advent of next generation sequencing technologies has enabled the high-throughput determination of the TCR sequence, allowing researchers to obtain the identity of TCRs and quantify the abundance and dynamics of T cell clones. Tracking clonal dynamics across different stages of disease and along the course of treatment have yielded valuable insights into T cell response against virus infection and cancer. Studies looking at SARS-CoV-2 reactive T cells have provided insights into repertoire changes along the course of repetitive infection or vaccination^8^, helping to identify protective components of adaptive immune system. In context of cancer, TCR tracing has helped to understand the mechanisms underlying immune checkpoint blockade^9–11^ and tumour-infiltrating lymphocyte (TIL) therapy^12^.

As with whole transcriptome profiling, TCR sequencing can be carried out at both bulk and single cell resolutions, with each methodology having its unique strengths and weaknesses. Although bulk TCR sequencing (TCR-seq)^13,14^ has the advantage of profiling the repertoire at a higher depth, allowing for the detection of rarer clones, it loses the *α*–*β* pairing information a key factor contributing to antigen recognition. On the other hand, single cell TCR sequencing (scTCR-seq), either droplet-based^15–18^, plate-based^19,20^ methodologies, or combinatorial-barcoding^21^, can preserve *α*–*β* pairing, and can be coupled to transcriptome profiling, allowing researchers to study repertoire in context of phenotypes^22,23^. However, single-cell methodologies are associated with several limitations, including a relatively high cost, low single cell capture rate, as well as transcript dropouts, leading to incomplete TCR recovery which are common in single-cell datasets^24,25^. Furthermore, the sensitivity of scTCR-seq is much lower than bulk TCR-seq, capturing 10^2^-10^3^ TCRs while bulk typically scales to 10^5^-10^6^ TCRs^22^.

When longitudinally tracking the dynamics of *α*–*β* paired clonotypes, repetitive scTCR-seq is often performed for samples from different timepoints, leading to significant cost. However, repetitive scTCR-seq to trace the expansion or contraction of recurring clones longitudinally results in an inefficient use of resources, by sequencing the same clones at more than one timepoints.

To overcome the limitations of existing TCR sequencing approaches, we developed ClonoDeconv, a recursive non-negative least square (NNLS) based method that leverages TCR pairing information from a single scTCR-seq reference sample to estimate paired *αβ* clonotype frequencies from bulk TCR-seq data. By enabling the sensitive detection of low-frequency paired clonotypes across longitudinal samples without repeated scTCR-seq and high sequencing-depth, ClonoDeconv provides a sensitive, cost effective and scalable pipeline for studying T cell clonal dynamics over time.

## Results

### Overview of ClonoDeconv

ClonoDeconv uses paired TCR clonotypes, to construct a reference matrix from scTCR-seq data to record known *α*–*β* pairings, resulting in different *α*–*β* paired clonotypes exhibiting distinct expression patterns of *α*/*β* chains. For cells expressing more than one *α* chain, the alternative pairings are weighted by their respective expression levels. Since individual *α*/*β* chain can pair with multiple distinct partners, clonotypes are frequently found to share chains with each other, leading to difficulties in estimating their individual frequencies. To address this, all *α*–*β* paired clonotypes in the reference matrix are first clustered into clonotype clusters/groups via graph-based or distance-based clustering methods (Methods). Clonotypes within a cluster have shared *α*/*β* chain, while different clusters are relatively distinct. In the first stage, the frequencies of clonotype clusters are estimated by non-negative least squares (NNLS). Subsequently, the clonotype clusters’ frequencies are further decomposed into individual clonotype frequencies with carefully subsetted bulk-TCR targets for each clonotype cluster that only include relevant chains to distinguish individual clonotypes apart (Figure 1). This generates estimated paired clonotype frequencies from the bulk TCR-seq data that can be utilised for further downstream investigation.

**Figure 1.**
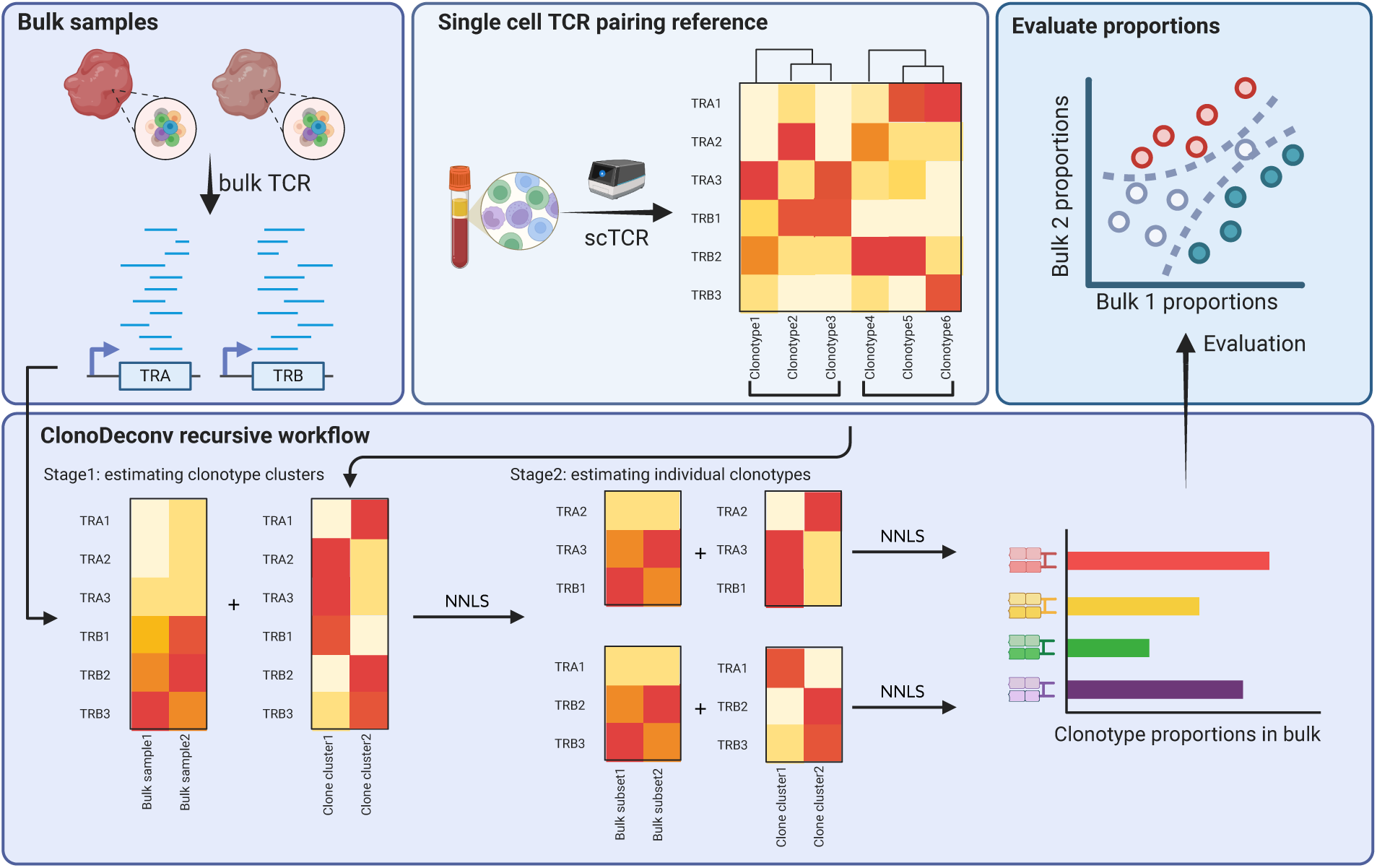
Illustration of ClonoDeconv workflow. In brief, ClonoDeconv takes in bulk TCR-seq samples as deconvolution targets, and constructs a pairing reference from an independent scTCR sample. All clonotypes in the reference are first grouped into clonotype clusters based on chain sharing. Frequencies of clonotype clusters are first estimated with NNLS. In the second stage, for each clonotype cluster, bulk target for the given cluster is compiled by subsetting chains in the complete bulk-TCR to only those that contribute to the distinction of clonotypes with the cluster, then frequencies of individual clonotypes can be derived from this subsetted deconvolution process. The derived *αβ* paired frequencies can be further used to study clonodynamics across bulk TCR-seq samples.

### Validating ClonoDeconv with simulated pseudo-bulk from 10x scTCR

To validate the feasibility of the method, two independent aliquots of the same melanoma metastatic lymph node single cell suspension were subjected to 10x scTCR-seq library construction and sequenced in parallel. One sample was kept as a scTCR-seq dataset and used as a reference in the deconvolution process, while the other sample was aggregated into a pseudobulk to simulate a bulk-TCR sample. This reference setup ensures a theoretically perfectly-matched TCR repertoire between the target pseudo-bulk and the reference, given they are from the same single cell suspension but still mimic a real scenario in which targeted bulk-TCR and the scTCR-seq reference are from different experiments (Figure 2a). The two scTCR runs (OX1a and OX1b) captured 2332 and 3156 unique *α*–*β* paired clonotypes respectively. As expected, the *α*–*β* V-gene pairing pattern was highly concordant across the two replicates (Figure 2b-d). Looking at the detected unique *α*/*β* chains, there was a significant overlap across both *α* and *β* compartments (Supplementary figure 1a, b), indicating a similar underlying clonotype distribution for the two datasets.

**Figure 2.**
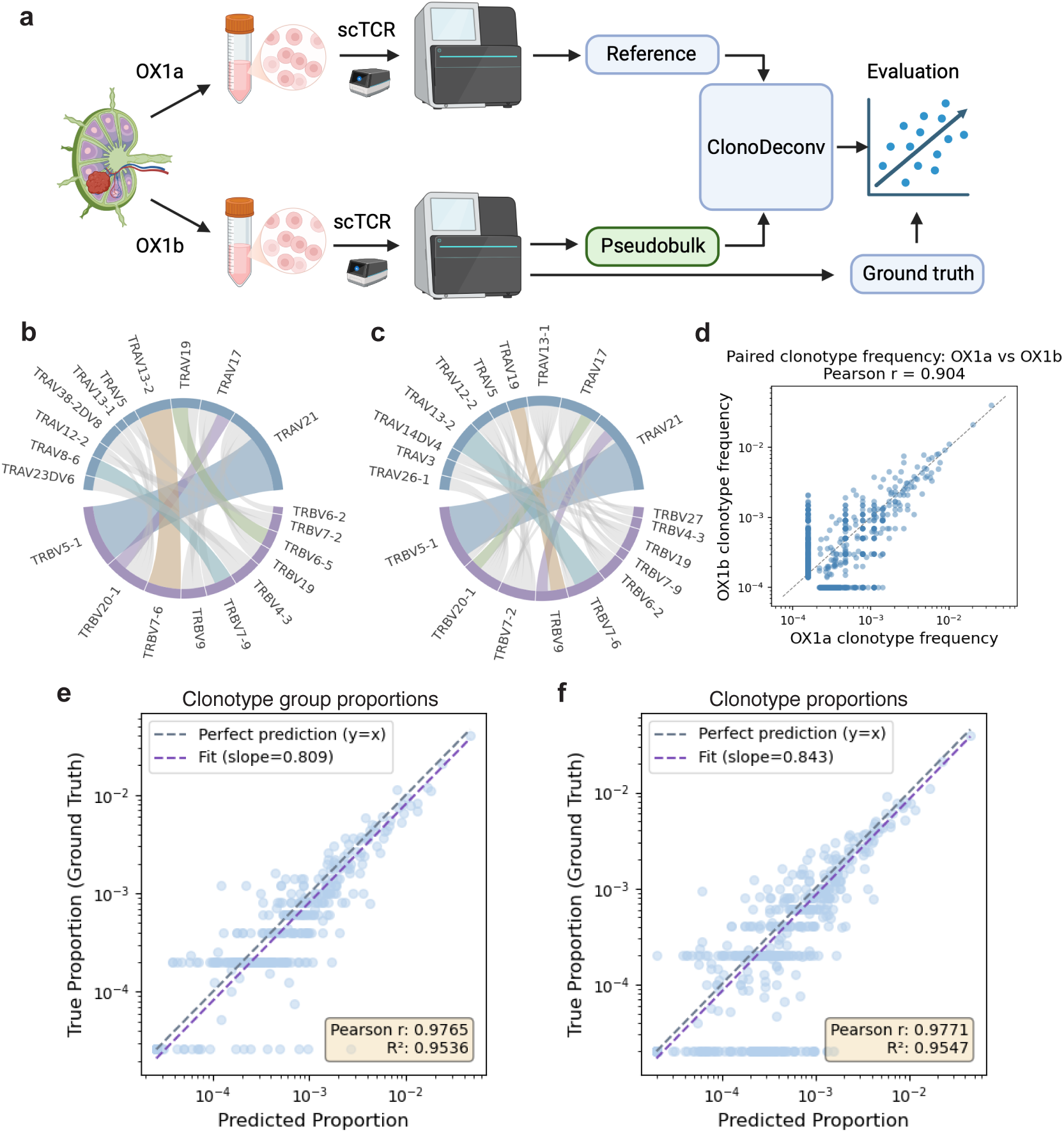
Validating ClonoDeconv with simulated pseudo-bulk from 10x scTCR-seq. **a**, Schematic of the experimental and analysis workflow; **b**, Circos plot showing top 10 *αβ* pairing in OX1a; **c**, Circos plot showing top 10 *αβ* pairing in OX1b; **d**, Paired clonotype frequency correlation between OX1a and OX1b scTCRseq. Each dot represents a unique clonotype and dotted line represents *y* = *x*, Pearson *r* = 0.904, *p <* 0.001; **e**, Scatter plot of deconvoluted clonotype group frequencies vs. ground truth frequencies in OX1b, blue dotted line shows *y* = *x*, purple dotted line is regression line, R^2^ is the coefficient of determination; Pearson *r* = 0.9765, *p <* 0.001; **f**, Scatter plot comparing deconvoluted clonotype frequencies of individual clonotypes vs. ground truth frequencies in OX1b, blue line shows *y* = *x*, purple dotted line is regression line, R^2^ is the coefficient of determination; Pearson *r* = 0.9771, *p* = 7.04 *×* 10*^−151^*.

Next, one scTCR-seq dataset (OX1b) was aggregated to simulate a bulk-TCR profile, while OX1a was used as the reference for deconvolution. After deconvolution, the estimated clonotype proportions from OX1b pseudo-bulk were compared to ground-truth proportions calculated from the OX1b scTCR-seq (Methods). ClonoDeconv first estimated the abundance of clonotype groups (Figure 2e, Supplementary figure 1c), then refined the analysis to individual *α*–*β* paired clonotypes (Figure 2f), achieving coefficient of determination R^2^ of 0.9536 and 0.9547 respectively. Moreover, ClonoDeconv was able to recapitulate 63% of the unique clones from OX1b pseudobulk using OX1a reference. At the individual clonotype level, compared to true TCR*αβ* frequencies, ClonoDeconv gave better overall frequencies when compared to a näıve one-stage NNLS (R^2^ = 0.9466, Supplementary figure 1d). Overall, these results suggest that ClonoDeconv is able to give reliable estimation of clonotype frequencies from pseudo-bulk data.

### Evaluating ClonoDeconv with real bulk-TCR dataset paired with SMARTer^®^ scTCR-seq

Having observed the optimal performance of ClonoDeconv in the pseudo-bulked dataset, we next tested its performance on a real bulk-TCR dataset. The bulk-TCR data was generated from a tumour-reactive CD8^+^ short-term T cell line from a breast cancer patient^26^, and paired SMARTer^®^ scTCR was generated from the same tissue. The *α*–*β* pairing information (without clonotype abundance) in the scTCR-seq dataset was used as the reference when deconvoluting the bulk TCR-seq dataset. After deconvolution, the inferred clonotype proportions were compared to the ground truth proportions derived by counting clonotypes in the scTCR-seq dataset to evaluate accuracy.

Limited by the scale of SMARTer^®^ scTCR-seq (152 cells with both successful *αβ* reconstruction), the single-cell dataset here only captures a subset of unique *α* chains and unique beta chains (Supplementary figure 2a) present in the bulk-TCR data that were generated from 1 million cells. However, the top 15 most abundant *α*/*β* chain pairings (Supplementary figure 2b) are captured by both bulk-TCR and SMARTer^®^ scTCR-seq (Figure 3a, b).

**Figure 3.**
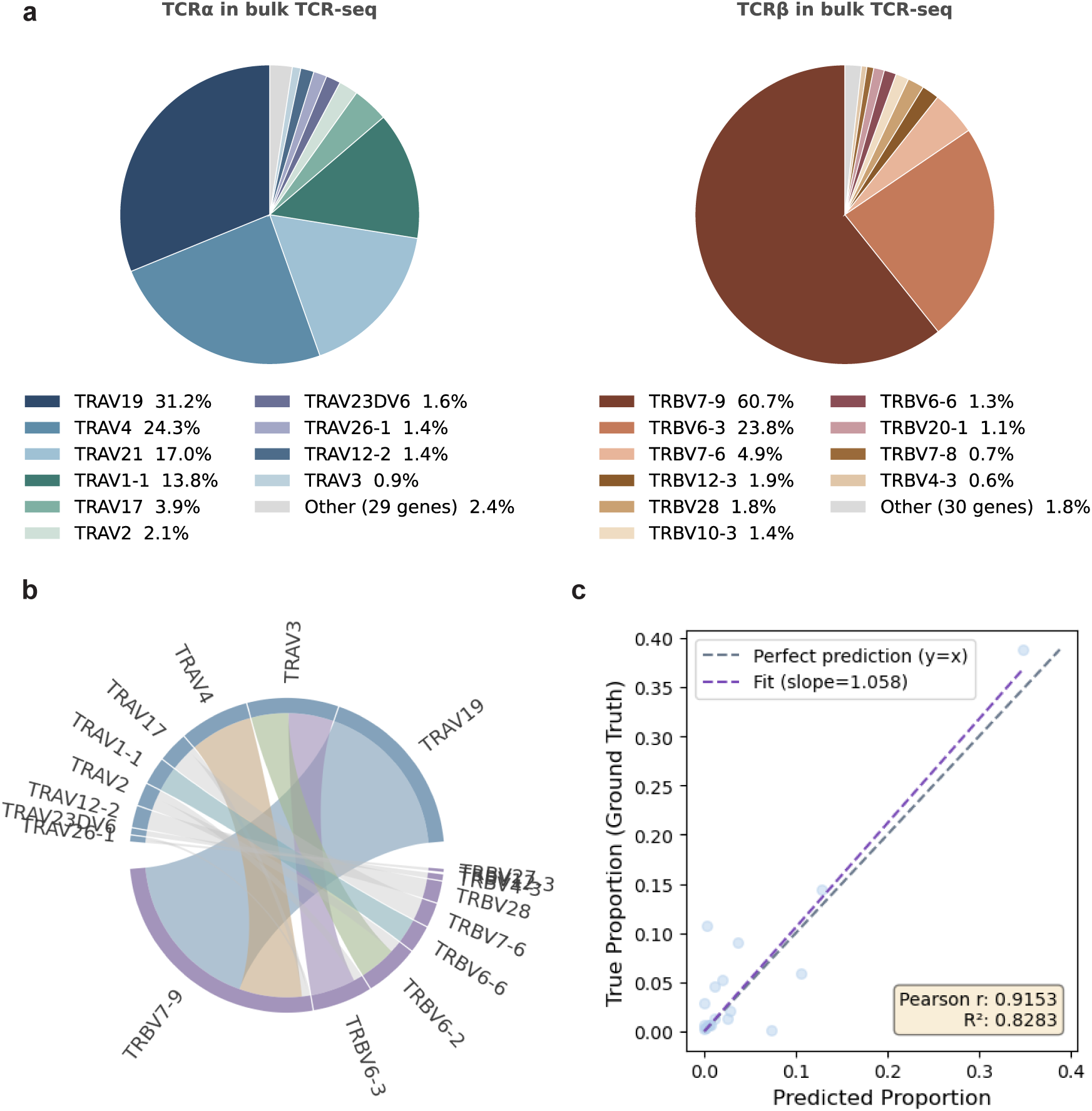
Evaluating ClonoDeconv with real bulk-TCR dataset paired with SMARTer^®^ scTCR-seq on tumor-specific CD8^+^ from breast cancer patient. **a**, Pie chart showing the frequencies of *α* and *β* gene usage in the bulk-TCR sample. **b**, Circos plot showing top 9 *αβ* pairings in SMARTer^®^ scTCR-seq data. **c**, Scatter plot comparing deconvoluted clonotype frequencies from bulk TCRseq v.s. frequencies in the scTCRseq dataset, blue line represents *y* = *x* and the purple line represent the fitted line, R^2^ is the coefficient of determination; Pearson *r* = 0.9153, *p* = 4.00 *×* 10*^−^*^08^.

Applying ClonoDeconv on the bulk-TCR data, all clonotypes were clustered into clonotype clusters and their proportions were estimated. ClonoDeconv identified the same abundant *α*–*β* pairings, and the estimated proportions correlated significantly with proportions derived from scTCR-seq (R^2^ = 0.8283) (Figure 3c). Furthermore, all of the identified clonotypes identified by the scTCRseq were recapitulated by ClonoDeconv from the bulk TCR-seq data from the same patient.

The above results highlight that ClonoDeconv is able to take TCR*α*/*β* data generated from different scTCR-seq platforms and could reliably estimate clonotype proportions from real bulk-TCR datasets, despite the difference in depth between the two techniques.

### Tracking *α*–*β* paired clonal dynamics in longitudinal bulk-TCR samples with ClonoDeconv

One of the primary usages for ClonoDeconv is to investigate clonal dynamics in longitudinal samples. In longitudinal experiments comprising samples from multiple timepoints/conditions from the same individual, a key question is the expansion or contraction of clonotypes across timepoints/conditions. We use ClonoDeconv to investigate whether we can estimate clonotype frequencies of overlapping *αβ* paired clonotypes in longitudinal samples using a scTCRseq reference dataset all generated from the same individual, to track clonal dynamics over time, with the aim of reducing the number of scTCR-seq experiments.

Here, ClonoDeconv was implemented using data from a publicly available dataset^27^ where PBMC samples were taken at three timepoints from an individual infected with severe acute respiratory syndrome coronavirus 2 (SARS-CoV-2). Baseline/tp1 sample was collected at 143 days before positive PCR test, acute/tp2 at 6 days after a PCR-positive test, and convalescent/tp3 at 29 days post-PCR positive test. The patient was seronegative for both SARS-CoV-2 and Epstein–Barr virus (EBV) at tp1, and later became seropositive for both viruses at the later timepoints. For each sample, parallel 5’ RACE bulk-TCR and 10x scTCR-seq for CD3+ T cells were performed. We used tp2 scTCR-seq dataset as the reference to derive *α*–*β* pairing and guide the deconvolution of the other two timepoints, as clones specific for both SARS-CoV-2 and EBV should be the most expanded at the acute timepoint (Figure 4a).

**Figure 4.**
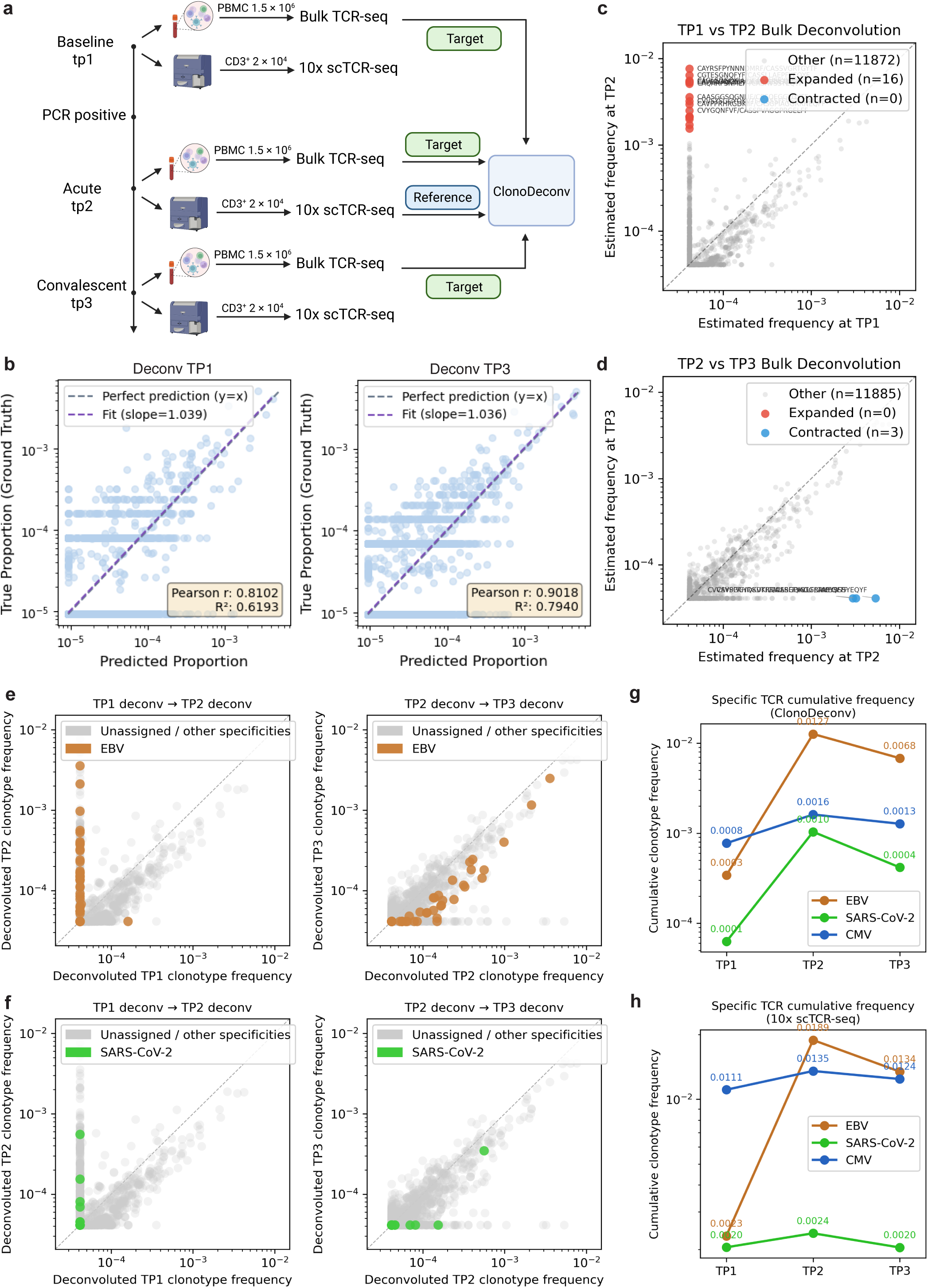
Tracking *αβ* paired clonal dynamics in longitudinal bulk-TCR samples with ClonoDeconv. **a**, Schematic of the experimental and analysis workflow, adapted from Pogorelyy et al., 2025^27^. **b**, Scatter plot comparing deconvoluted clonotype frequencies from the bulk-TCR sample v.s. ground truth clonotype frequencies in the scTCRseq dataset. Left panel=acute/tp1, Pearson *r* = 0.8102, *p* = 5.20 *×* 10*^−^*^109^; right panel=convalescent/tp3, Pearson *r* = 0.9018, *p* = 6.17 *×* 10*^−^*^234^. Blue line represents *y* = *x* and the purple line represent the fitted line, R^2^ is the coefficient of determination. **c**, **d**, Identification of significantly expanded (red) and contracted (blue) clones between pairwise timepoints (**c**, tp1 to tp2; **d**, tp2 to tp3). Significance given by log2 fold change *>*1 or *<* -1, and FDR *<* 0.05 (Fisher exact test with BH multiple comparison correction). All other non-significant clonotypes are in grey. **e**, Clonal dynamics of EBV-specific clones (n = 85, yellow) between pairwise timepoints (left tp1-tp2, right tp2-tp3). **f**, Clonal dynamics of SARS-CoV-2 specific clones (n = 27, green) on pairwise timepoints (left tp1-tp2, right tp2-tp3). **g**, Cumulative frequencies of EBV-specific (orange), SARS-CoV-2-specific (green) and CMV-specific (blue) clonotypes across timepoints, as estimated with ClonoDeconv. **h**, Cumulative frequencies of EBV-specific (orange), SARS-CoV-2-specific (green) and CMV-specific (blue) clonotypes across timepoints, as present in the 10x scTCR-seq dataset.

Single cell results consistently captured less unique *α*/*β* chains than bulk-TCR (Supplementary figure 3a-c, g). Despite this, there remains a good overlap between bulk-TCR and scTCR-seq from the same samples, with 30-36% of TRA chains and 21-23% TRB chains identified by scTCRseq also found in the corresponding bulk TCRseq sample at the three timepoints, particularly evident for the top 10% abundant chains (Supplementary figure 3d-f, h). Furthermore, we observed around 47% TCR*α* and 31% TCR*β* chains from tp1/tp3 overlapped with tp2 bulk (Supplementary figure 4a, b).

Using ClonoDeconv, we identified 3985 and 4882 paired clonotypes at differing frequencies at tp1 and tp3 respectively. The deconvoluted clonotype frequencies at tp1 and tp3 were compared to the true frequencies derived from scTCR-seq at the same timepoints to evaluate the deconvolution accuracy. ClonoDeconv achieved R^2^ of 0.63 and 0.80 for tp1 and tp3 respectively (Figure 4b). Comparing the deconvoluted clonotype frequencies at tp1 and tp3 to tp2, ClonoDeconv identified 16 clonotypes that significantly expanded, as well as 2 contracted clonotypes between tp2 and tp3 (Figure 4c, d, Supplementary table). The clones identified as significantly expanded/contracted using ClonoDeconv had 72% of overlap with those identified as expanded/contracted between timepoints by scTCR-seq of all timepoints (14 expanded and 2 contracted clonotypes from tp1 to tp2; 2 contracted clonotypes from tp2 to tp3, Supplementary figure 5a and Supplementary table). These results suggest the feasability of the ClonoDeconv workflow as a cheaper and more scalable alternative to carrying out scTCR-seq at multiple timepoints.

### Tracking antigen-specific clonal dynamics with ClonoDeconv

Having shown that ClonoDeconv is able to recapitulate clonal dynamics, we next examined whether clonal dynamics of rarer antigen-specific populations are preserved and can be investigated. We identified clonotypes with known antigen-specificities documented in the VDJdb^28^. We annotated the clonotypes by mapping to paired TCR entries in the VDJdb with EBV/SARS-CoV-2 specificities and the same HLA restrictions as the donor, and tracked these clones across disease progression using the inferred ClonoDeconv frequencies.

We observed a clear expansion of EBV-specific clones from baseline to acute samples was observed (Figure 4e). Furthermore, the cumulative frequency of clones recognizing EBV epitopes rose from tp1 to tp2 and reduced by 46% at tp3. For SARS-CoV-2 specific clones, a similar pattern of clonal expansion followed by contraction was observed (Figure 4f). Similar to the frequency of EBV clonotypes, the cumulative frequency of SARS-CoV-2 specific clones increased from tp1 to tp2 and reduced 60% at tp3 (Figure 4g). In contrast, the cumulative frequencies of clones with inferred CMV-specificity remained relatively stable across timepoints. This was consistent with the hypothesis that the patient with mild disease was highly unlikely to be undergoing CMV reactivation during the sampling window as CMV reactivation mostly happens in critical patients especially with immunosuppression^29^.

Investigating the clonal dynamics of the same EBV/SARS-CoV-2/CMV clones with frequencies calculated from the 10x scTCR-seq data revealed largely similar trends (Supplementary figure 5b, c and Figure 4h). However, the expansion and contraction of SARS-CoV-2 specific clones were not observed in the scTCR-seq data, likely due to the lower number of cells captured in the 10x scTCR-seq dataset. These changes in the EBV/SARS-CoV-2-specific clonal dynamics observed by ClonoDeconv indicate the occurrence of the two infections within the sampling window, corresponding with the change of serostatus of the patient^27^. Overall, ClonoDeconv was able to identify changes in clonotypes frequencies, both antigen-specific and total repertoire dynamics between multiple longitudinal samples, providing insights into the dynamics of the less abundant SARS-CoV-2-specific clones that were not observed in the 10x scTCR-seq dataset.

**Figure 5.**
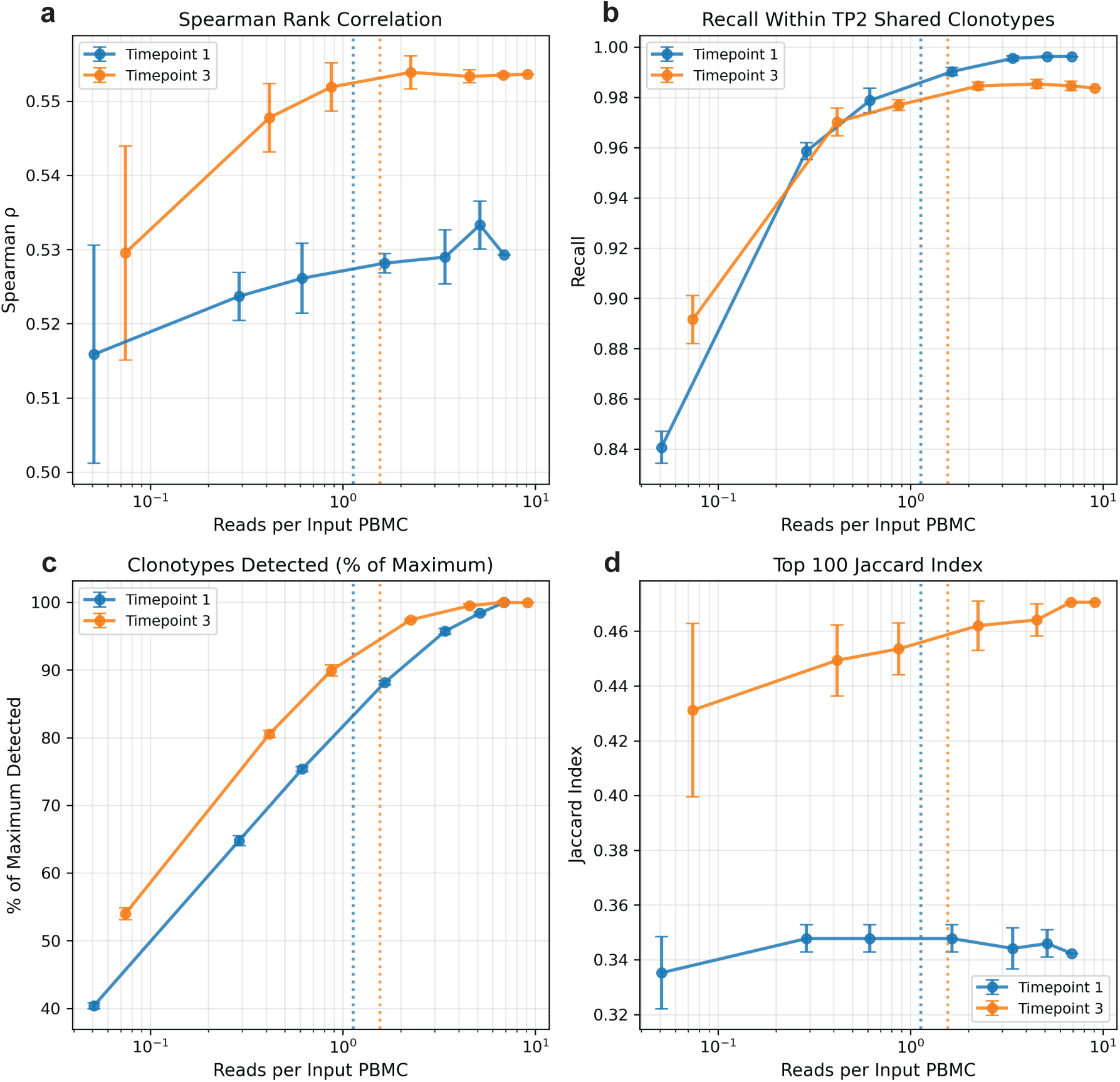
Robustness against sequencing depth. **a**, Spearman correlation between ClonoDeconv estimated frequencies and 10x scTCRseq ground truth frequencies for overlapping clonotypes across timepoints, each point represents one downsampling fraction of one sample, error bar denotes the standard error across replicates. **b**, Fraction of clonotypes detected (estimated frequency *>* 0) amongst all clonotypes shared by target bulk (blue: tp1, yellow: tp3) and tp2 reference. **c**, Fraction of clonotypes detected (estimated frequency *>* 0) among all clonotypes that can be detected at full sequencing depth. **d**, Jaccard index between estimated top 100 clonotypes in each downsampling fraction and top 100 clonotypes from 10x scTCRseq dataset of the corresponding timepoint. Across all four panels, the dotted lines indicate the sequencing depth corresponding to the midpoint between two downsampling fractions (10%-25% of the original sequencing depth), where a noticeable decrease in recall among tp2 shared clones occurs.

### Robustness against bulk-TCR sequencing depth

As the number of unique clonotypes reconstructed by bulk-TCR sequencing is generally affected by overall sequencing depth^30^, ClonoDeconv’s performance against serial sequencing depth was evaluated. Using the above longitudinal dataset, tp1 and tp3 bulk-TCR raw FASTQ files were downsampled randomly to a series of fractions (1%, 5%, 10%, 25%, 50%, and 75%) to simulate different sequencing depth. For each downsampling fraction of the same sample, 5 replicates of downsampling were performed to evaluate stability of estimates.

Correlating the clonotype frequencies derived from the ClonoDeconv pipeline with frequencies given by the scTCR-seq dataset at each corresponding timepoint, the Spearman correlation remained largely stable above 10% of the original sequencing depth (Figure 5a). Looking at the fraction of clonotypes that have estimated frequencies above 0 among all clonotypes shared by target bulk-TCR (tp1/tp3) and reference (tp2) (Figure 5b), and the fraction of clonotypes detected among all clonotypes that could possibly be detected with full depth (Figure 5c), both metrics remain stable and don’t show a noticeable decrease until subsetting to 25% of the original read depth. The Jaccard index between the top 100 estimated clonotypes and the top 100 clonotypes in scTCR-seq data from corresponding timepoints stayed mostly stable (Figure 5d).

Across all metrics of deconvolution accuracy and sensitivity, the results remained largely concordant with the full sequencing depth until downsampled to around 25% of original depth, where a noticeable decrease of performance started to be observed. This corresponds to around 1 sequencing read per PBMC input into bulk TCR-seq, demonstrating a low requirement of sequencing depth.

## Discussion

In this study, we have demonstrated that the ClonoDeconv pipeline is able to accurately predict *α*-*β* paired clonotype frequencies from bulk-TCR datasets for clonotypes that overlap with a single separate scTCR-seq reference, greatly reducing the overall experimental cost for longitudinal studies, making repertoire evaluation scalable to more samples. The feasibility of this approach has been validated across T cell responses against both cancer and virus infection. Furthermore, the two-stage recursive NNLS method utilized in ClonoDeconv gives more accurate results than naive one-stage NNLS across all of our tests. ClonoDeconv works agnostically of the methodology used to generate both the single-cell TCR reference and the bulk TCR datasets, providing flexibility to researchers. Finally, ClonoDeconv was robust against various sequencing depth in bulk TCR-seq.

Compared to scTCR-seq technologies, ClonoDeconv has the advantage of capturing rarer clonotypes due to the broad coverage of bulk TCR-seq. The throughput of the state-of-the-art scTCR-seq methods (e.g. 10x Genomics Immune profiling, BD Rhapsody TCR assays) is typically around 10^4^ cells, which represents a small proportion of all T cells in the human body, which are predicted to be around 10^12^ cells and 10^8^ clonotypes^31^. As such, the lowest frequency such a scTCR-seq experiment can robustly provide is approximately 10^-4^, with over 99% of clonotypes below 10^-4^ ^32^, making estimation of these clonotypes inaccurate^33^. Moreover, if sampling the clonotypes randomly from the whole sample, for rarer clones, the observed clone frequency in a scTCR-seq experiment follows an approximate Poisson distribution^34^, with a coefficient of variation inversely correlated with expected frequency and sampling size. Therefore, rarer clones (with low expected frequency) will naturally have an extremely unstable frequency estimation given by scTCR-seq with a limited sampling size, due to random sampling effects, similar to what has been observed in RNA-seq modeling studies for lowly-expressed genes^35,36^. On the other hand, bulk-TCR sequencing is able to sample clonotypes from a larger cell population, and as such, is able to better detect less abundant clones, providing relatively more stable frequencies, albeit without *αβ* pairing information. By extracting only the pairing information from scTCR-seq (which *α* chain pairs with which *β* chains), clonotypes frequencies can be estimated by ClonoDeconv from bulk TCR-seq, therefore providing a more stable estimation with a lower detection limit compared to traditional scTCR-seq methods. This superiority over scTCR-seq has been observed when tracking SARS-CoV-2 specific clones, where the ClonoDeconv recapitualated the expansion and contraction of SARS-CoV-2 clones, while the scTCR-seq method failed to do so.

Another method for deriving *αβ* paired clonotype frequencies is through combinatorial TCR pairing strategies^27,37–39^. This approach infers pairing from multiple replicates of bulk TCR-seq. Compared to combinatorial approaches, once a scTCR-seq reference has been generated, ClonoDeconv only requires one bulk TCR-seq dataset per sample, which can be generated using as little as 1ng of total RNA, allowing potentially more samples to be evaluated at the same time. This low sample input requirement per sample to be investigated for bulk TCR-seq makes it more suitable when the starting material is limited, as can be the case for many clinical samples. For example when tracking low frequencies of antigen-specific T cells or studying T cell clones from non-lymphoid tissues, where hundreds of bulk TCR-seq replicates may not be feasible. Furthermore, combinatorial approaches are unable to estimate the frequency of clones with low frequencies as they rarely appear in more than one replicate^27,39^. ClonoDeconv, however, has the ability to estimate such low numbers from bulk TCR-seq, albeit with the limitation that these rarer clones have to be covered in bulk TCR-seq of enough depth, and the TCRs are found in scTCR-seq reference and as such the pairings could be empirically determined.

ClonoDeconv could be applied to investigate T cell clonal dynamics across conditions and timepoints. This is of particular use in cancer and investigating cancer immunotherapy. When studying the T cell response to immunotherapy, using pre-therapy tumor-infiltrating and blood T cells as scTCR-seq reference, it is possible to investigate the expansion or contraction of T cell clones responding to therapy, both from within-tumour precursors^9^ and blood precursors^10,11^. Therefore, by carrying out bulk TCR-seq on post-therapy samples, is can be possible to investigate therapy-responsive clones that may play a role in the response. In cases when antigen-specificity can be surveyed, ClonoDeconv can also be applied to a more restricted repertoire, selected for antigen-specificity, as demonstrated for tumour-reactive cells in breast cancer. This also allows for the examination of antigen-specific clonal dynamics along the course of infection^40,41^ or vaccination^42,43^, where antigen-specific T cell response is more conventional surveyed.

Beyond TCR repertoires, ClonoDeconv may also facilitate longitudinal tracking of antigen-specific B cell responses. By leveraging paired single-cell BCR sequencing as a reference, the framework could enable estimation of paired heavy- and light-chain clonotype frequencies from bulk BCR sequencing datasets collected across multiple time points. Such an approach could prove valuable for monitoring vaccine-induced clonal expansion, affinity maturation and contraction of antigen-specific B cell populations, as well as characterizing humoral immune dynamics during infection, autoimmune disease or cancer. Integrating the scalability of bulk BCR sequencing with paired-chain information from single-cell datasets may therefore provide a cost-effective strategy for studying antibody repertoire evolution and identifying clinically relevant B cell clonotypes associated with protective immunity, disease progression or therapeutic response.

Despite the advantages, one important limitation of our method is the inability to estimate frequencies of previously unobserved pairings. Only *α*-*β* paired clonotypes that have been observed in the selected scTCR-seq reference can be estimated in subsequent bulk TCR-seq samples. Therefore, Clonotype overlap between the scTCR-seq reference and targeted bulk TCR-seq should be inspected before running the estimation to ensure adequate clonotype overlap, as well as careful selection of the reference sample, such that the scTCR-seq reference should capture the majority of the biologically relevant clones, resulting in the overlap between the scTCR-seq and bulk TCR-seq datasets. Furthermore, the frequency estimation of ClonoDeconv may also benefit from careful selection of the target bulk. For example, estimating within selected compartments (e.g. T cell populations sorted based on their phenotype or antigen-specificity) could greatly reduce noise from irrelevant clones in whole PBMC that may account for a large proportion of the T cell population in the whole PBMC sample. For example, when investigating CD8+ clonal dynamics, selection of CD8+ T cells for targeted bulk-TCR sequencing will reduce the noise from CD4+ clonotypes.

Overall, ClonoDeconv offers an easy-to-implement, scalable and cost-effective approach to estimate *α*-*β* paired clonotype frequencies from bulk-TCRseq datasets using a paired scTCR-seq dataset as a reference. We anticipate this method could be used in tracking clonotype frequencies across timepoints, across tissue compartments and between conditions, highlighting biologically relevant T cell clonotypes for further mechanistic investigation and personalized therapy development.

## Methods

### Lymph node tissue processing

Lymph node biopsies were manually dissociated and minced using a scalpel on a sterile petri dish on ice. Dissociated tissue was rinsed with cold PBS and transferred to a falcon tube. Sample was centrifuged for 5 minutes at 1500 rpm at 4*^◦^*C. Supernatant was discarded and pellet resuspend in 5 ml cold PBS+0.4% BSA. Resuspended cells/tissue was passed through a 70 *µ*m strainer into a sterile 50 mL tube and any remaining tissue was gently pressed onto the strainer to pass any remaining cells. Strainer was rinsed twice with 5 ml cold PBS+0.4% BSA. Cells were centrifuged for 5 minutes at 1500 rpm at 4*^◦^*C, supernatant discarded and pellet resuspend in 5 ml cold PBS+0.4% BSA.

### 10X 5’ Immune Profiling with Feature Barcodes

250,000 cells were washed with PBS+0.4% BSA and resuspended in 47.5 *µ*l of PBS+0.4%BSA. 2.5 *µ*l Human TruStain FcX (Fc receptor blocking solution, Biolegend 422301) was added and cells incubated at 4*^◦^*C for 10 min. Cells were washed twice with PBS+0.4%BSA and resuspended in 50ul PBS+0.4%BSA. 50 *µ*l of reconstituted TotalSeq-C Human Universal Cocktail (prepared following manufacturer’s instructions) was added and incubated at 4*^◦^*C for 30 mins. Cells were washed twice with PBS+0.4%BSA and finally resuspended in PBS+0.4%BSA at 2000 cells/*µ*l. 50,000 cells were loaded into two independent lanes of a 10X 5’ Immune profiling chip and processed following the manufacturer’s instructions. cDNA was amplified using 13 PCR cycles and gene expression, TCR and feature barcode libraries generated. Libraries were pooled at a ration of 5:1:3 (gene expression:TCR:feature barcode) and sequenced on a NovaSeq X (Novogene UK) for a total of 70,000 reads per cell.

### Obtaining tumour-reactive T cells from breast cancer patients for SMARTer^®^ scTCR and bulk-TCR

Sample processing and sequencing were carried out as previously published^26^. In brief, T cells were isolated from patient Peripheral blood mononuclear cells (PBMCs) using Pan T Cell Isolation Kit (Miltenyi Biotec), following the manufacturer’s instructions. Then, T cells were labelled with CFSE (Thermo Fisher Scientific) and cocultured with autologous tumour lysate loaded dendritic cells (DCs). Tumour-reactive cells were harvested at day8 and sorted for proliferating CD8^+^ cells (CD8+CFSElowCD25+). Short-term bulk T cell lines were generated by sorting proliferating CD8^+^ in the same tube and expanded using irradiated allogeneic PBMCs with PHA as previously described^44^. At day 14 of expansion, 10^6^ T cells from the short-term line were subjected to bulk-TCR sequencing. while single T cells were sorted into each well of 96 plate for SMARTer^®^ scTCR sequencing.

### SMARTer^®^ single-cell TCR sequencing

Tumour-reactive CD8 T cells isolated from the proliferation assay were expanded and individually sorted into 96-well plates containing 5.75 *µ*L of lysis buffer, composed of 1× lysis buffer supplemented with RNase inhibitor. Plates were stored at -80*^◦^*C until downstream processing. Single-cell TCR profiling was performed using the SMARTer^®^ Human scTCR *α*/*β* Profiling Kit (Takara) according to the manufacturer’s protocol. Briefly, lysed samples were transferred into fresh 96-well plates. 4 wells of Jurkat RNA and 4 wells of water were included as positive controls and negative controls respectively. First-strand cDNA synthesis was then performed using row-specific SMARTer^®^ Index Oligos, followed by cDNA amplification. Samples from each column were pooled, and the resulting cDNA was purified with SPRIselect beads (Beckman Coulter).

Nested PCR amplification was carried out using TCR-specific primers together with indexed primers to enable sample multiplexing. The amplified libraries were purified using SPRIselect beads and quantified with High Sensitivity D5000 reagents on an Agilent TapeStation. Equimolar library pools were prepared and sequenced on an Illumina MiSeq platform using the 600-cycle MiSeq Reagent Kit v3, generating paired-end 2 × 300 bp reads. BCL files were demultiplexed and converted to FASTQ format using bcl2fastq v.2.20.0.422 (Illumina). Cells were demultiplexed using the SMARTer Human scTCR Demultiplexer (Takara) and TCR clonotypes were extracted using MiXCR v.3.0.13^45^.

### Bulk TCR sequencing

Cells were washed with phosphate-buffered saline (PBS), after which total RNA was isolated using RNeasy Plus Mini Kit (QIAGEN). RNA concentration was measured using a NanoDrop spectrophotometer (Thermo Fisher Scientific).

RNA extracted from T-cell lines was used to prepare full-length TCR repertoire libraries for Illumina sequencing with the SMARTer Human TCR *α*/*β* Profiling Kit v2 (Takara), according to the manufacturer’s protocol and as previously described^8^. Briefly, cDNA synthesis was performed using primers containing unique molecular identifiers (UMIs). cDNA was then subsequently amplified through two rounds of PCR with nested primers. The resulting PCR products were purified using SPRIselect beads (Beckman Coulter) and eluted in EB buffer (QIAGEN).

Library concentration and quality were assessed on a 4200 TapeStation system using D5000 reagents (Agilent). Sequencing was carried out on an Illumina MiSeq platform using the MiSeq Reagent Kit v3, 600 cycles, with MiSeq Control Software v2.6.2.1. BCL files were demultiplexed and converted to FASTQ format using bcl2fastq v.2.20.0.422 (Illumina). TCR clones were extracted using MiXCR v.3.0.13^45^.

### Overview of ClonoDeconv workflow

ClonoDeconv is a Python package for estimating clonotype-level proportions from bulk TCR repertoire sequencing data, using paired single-cell TCR sequencing (scTCR-seq) as a reference. The workflow proceeds in four steps: (i) preprocessing of scTCR-seq data into a standardised chain-level format; (ii) construction of a reference matrix in which rows represent unique *α* or *β* chains and columns represent paired clonotypes; (iii) graph-based clustering of clonotypes that share one or more chains, and reconstruction of a grouped reference matrix; and (iv) two-stage recursive non-negative least squares (NNLS) deconvolution to estimate clonotype proportions in the bulk sample.

Because the bulk TCR-seq dataset routinely contains clonotypes absent from the single-cell reference, ClonoDeconv calculates a *coverage factor f*_cov_ — the fraction of total bulk counts attributable to chains present in the reference — and uses it to anchor the deconvolution result so that the sum of estimated proportions does not exceed one. This prevents systematic over-estimation of clonotype frequencies when reference coverage is partial.

The two-stage deconvolution first estimates proportions at the level of clonotype groups (clusters):

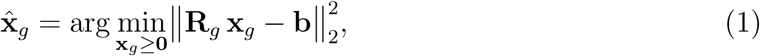

where **R***_g_* is the grouped reference matrix and **b** is the bulk count vector. A refined second-stage NNLS is then applied independently within each cluster to resolve individual clonotype proportions; the optimisation target for each cluster is constructed to incorporate only the expression signal attributable to that cluster, while excluding chains that do not distinguish its members (see Supplementary Methods for full equations).

### Data preprocessing

#### 10x **Genomics Cell Ranger VDJ output**

FASTQ files were processed using Cellranger v8.0.1 using `cellranger multi` to process gene expression, VDJ and feature barcode FASTQ files simultaneously. Raw output from Cellranger was filtered to retain only high-quality, full-length and productive contigs. TRA and TRB chains were separated by the chain column. Compatible with cells with more than one TRA chains detected, each unique *α* chain in the same cell was weighted by within-cell chain fractions. The within-cell chain fractions were computed as the UMI count of each chain divided by the total UMI count across all chains of the same type in that cell. An unique single chain was defined as the concatenation of the V-gene name and the CDR3 amino acid sequence. UMI-based 10x counts were used directly without batch normalisation, as UMI counts are inherently corrected for amplification bias.

#### SMARTer^®^ MiXCR output

BCL files were demultiplexed using bcl2fastq, and the resulting FASTQ files were further processed with the SMARTer Human scTCR Demultiplexer (Takara) to separate pooled single-cell samples within each FASTQ file. The first 41 bases of read 2 were trimmed using `cutadapt` to remove the SMART sequence and inline indices.

Clonotypes for individual cells were identified using MiXCR v3.0.13^45^ with the `analyse amplicon` preset. Plate-specific read-count thresholds were calculated from the negative-control wells as:

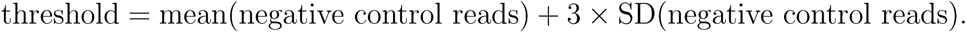

Only cells with clonotype read counts above the corresponding threshold were retained for downstream analysis. Cells containing clonotypes detected in the Jurkat positive-control wells were excluded, as these were considered potential cross-contamination events.

In the filtered list, single chain were defined identically to the 10x pipeline. To correct for differences in sequencing depth across multiplexed sequencing plate columns, read counts were batch-normalised: for each plate column, the counts of all chains were divided by the total reads in that batch and multiplied by the number of cells in the batch, yielding within-batch normalised counts with a common scale.

#### Paired clonotype definition

Only cells with at least one *α* and one *β* chain passing quality filters were retained. For each cell, the highest-UMI (or highest-read-count for SMARTer^®^ scTCR-seq) *β* chain was selected. All *α* chains with a within-cell frequency above a minimum threshold (default 0.2) were retained, allowing dual-alpha cells. Cells with multiple *α* chains were split into multiple clonotype records, weighted by relative alpha-chain expression (UMI or read count ratio), so that each cell’s total contribution summed to one. A paired clonotype was defined as the concatenated string “V*α* CDR3*α*/V*β* CDR3*β*”.

### Simulating pseudo-bulk TCR profiles from scTCR-seq datasets

Pseudo-bulk TCR profiles were generated from the scTCR-seq dataset as an in-silico analogue of bulk repertoire sequencing, used for method validation. For each unique *α* or *β* chain in the reference, counts (normalised for SMARTer^®^ scTCR-seq, UMI-based for 10x) were summed across all cells. The resulting vector was indexed on the same chain identifiers as the reference matrix, providing a bulk dataset in which the ground-truth clonotype proportions are known exactly from the single-cell data.

### Evaluating performance on pseudo-bulk datasets

Deconvolution accuracy was evaluated using pseudo-bulk profiles derived from 10x VDJ data. Two single-cell samples from the same donor (OX1a and OX1b) were used: the reference matrix was constructed from one sample, and the pseudo-bulk was derived from the other, simulating the cross-sample deconvolution scenario. Single-cell clonotype proportions were computed from the single-cell data, accounting for dual-alpha cells by distributing each cell’s contribution in proportion to alpha-chain UMI counts. Estimated proportions from the two-stage NNLS were scaled by the coverage factor so that their sum equals *f*_cov_, matching the scale of the underlying ground-truth proportions. Performance was quantified by Pearson correlation coefficient *r* and the coefficient of determination *R*^2^.

Because scTCR-derived frequencies are often inaccurate for rare clones, the evaluation metrics were calculated for confident clonotypes present in sufficient frequencies in the scTCR ground truth among all clonotypes that have a deconvoluted frequency. The frequency threshold was set as the lowest frequency scTCR can give (only one cell detected), *N_scT CR_* is the number of cells in the scTCR-seq ground truth.

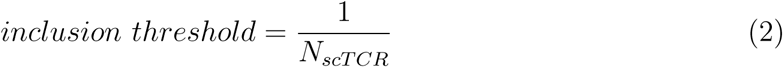

### Evaluating performance on real bulk-TCR data with paired SMARTer^®^ scTCR-seq reference

Validation on real bulk data was performed using a sample-matched bulk TCR-seq and SMARTer^®^ scTCR-seq dataset from a single patient (Br14). The scTCR-seq data provided the reference matrix after batch normalisation, and the corresponding bulk TCR-seq file (MiXCR-processed clonotype counts) served as the deconvolution target. Performance was assessed by correlating the deconvolved clonotype proportions against the single-cell proportions.

### Analyzing clonal dynamics on longitudinal samples with 10x VDJ

Longitudinal scTCR-seq and bulk TCR-seq data from the COVID patient^27^ was downloaded from Zenodo at https://doi.org/10.5281/zenodo.14010377.

To characterise clonal expansion and contraction across time points directly from 10x scTCR-seq data, clonotype frequencies were computed independently at each time point using the same preprocessing pipeline as ClonoDeconv. Specifically, Cell Ranger VDJ output was filtered as for melanoma data and clonotypes were defined as described in data preprocessing; dual-alpha cell weights were applied, and frequency was defined as the sum of weighted cell counts belonging to the given clonotype divided by the total number of paired cells at that time point (12446 cells at tp1, 11882 cells at tp2 and 14451 cells at tp3).

Differential abundance was tested for every clonotype observed at either time point using a two-sided Fisher’s exact test with Benjamini–Hochberg (BH) false-discovery rate (FDR) correction applied across all tests. A pseudocount of 0.5/ max(*N*_1_*, N*_2_) was added to both frequencies before computing log_2_ fold change (log_2_FC) to avoid undefined values for clonotypes absent from one time point. Clonotypes were classified as *expanded* if log_2_FC *>* 1 and *p*_adj_ *<* 0.05 (more than two-fold increase), and as *contracted* if log_2_FC *< −*1 and *p*_adj_ *<* 0.05.

### Analyzing clonal dynamics on longitudinal samples with ClonoDeconv

In the intended scenario, scTCR-seq was only carried out only at tp2, and bulk TCR-seq was obtained for all timepoints. At time points for which only bulk TCR-seq data were available, ClonoDeconv was applied to estimate clonotype-level frequencies using the 10x scTCR-seq dataset from time point 2 as the reference. The reference was built from tp2 10x scTCR-seq, and bulk TCR-seq at all timepoints were deconvoluted using the two-stage NNLS procedure. The resulting tp1-tp3 deconvoluted frequencies were compared to tp2 deconvolved frequencies to identify clones that changed in abundance.

Because the deconvoluted tp1 (or tp3) values are continuous estimates rather than integer cell counts, statistical testing was performed using a modified Fisher’s exact test in which the deconvoluted frequencies were converted to an equivalent count at tp2 cell count by multiplying the frequencies to tp2 cell counts. This is because the dataset used PBMC for bulk TCR-seq so the true T cell counts in bulk are not available. In practices where T cell counts in bulk as obtainable, clone counts can simply be derived by multiplying estimated frequencies by the total T cell counts. BH FDR correction and log_2_FC thresholds were applied as above.

### Identifying virus-specific clonotypes

Antigen specificity was assigned by co-clustering experimental paired *αβ* clonotypes with TCRs of known specificity from the VDJdb database^28^, downloaded from the release dated 29 December 2025.

VDJdb records were filtered to retain only entries satisfying all of the following criteria: (i) both *α* and *β* CDR3 sequences are reported; (ii) the TCR is human (*Homo sapiens*); (iii) the restriction is MHC class I; (iv) the restricting MHC allele matches at least one of the donor’s known HLA alleles (A*02, A*03, B*07, B*08, C*07). Only HLA class I were reported in the original study so class II alleles were not analyzed here; (v) the antigen species is SARS-CoV-2, CMV, or EBV; and (vi) the epitope is not one of two immunodominant influenza or CMV epitopes (GILGFVFTL, KLGGALQAK) that were excluded due to over-representation in public databases^27,46^.

Experimental paired clonotypes from all longitudinal comparisons were pooled, deduplicated, and combined with the filtered VDJdb TCRs. Pairwise TCR distances were computed using the TCRdist3 algorithm^47^ on the combined set with sparse computation (combined *α*+*β* TCRdist threshold of 90). Edges were drawn between pairs with a combined distance below 90, forming a similarity graph. To remove hub nodes that spuriously link otherwise unrelated clusters, the top 1% of nodes by betweenness centrality (approximated with *k* = 500 random pivots) were removed. Connected components of the pruned graph were evaluated for VDJdb enrichment: a component was assigned the specificity of an epitope if more than 40% of its nodes corresponded to VDJdb TCRs recognising that epitope. Experimental clonotypes belonging to an assigned component inherited that specificity label.

### Sequencing depth robustness analysis

To evaluate how deconvolution accuracy depends on sequencing depth, we performed a downsampling experiment on the bulk TCR-seq libraries from the longitudinal samples. For each timepoint (tp1, acute; tp3, convalescent), we generated downsampled replicates at six nominal depth fractions: 1%, 5%, 10%, 25%, 50%, and 75% of the original read count with Seqtk, plus the undownsampled (100%) library, yielding 5 independent replicates per fraction–timepoint combination.

TCRs from each downsampled library were reconstructed using MiXCR, and successfully reconstructed samples were further processed through the full ClonoDeconv two-stage NNLS pipeline. The tp2 (acute) 10x scTCR-seq data served as the paired-clonotype reference for all runs; the reference and its graph-based cluster structure were constructed once and reused across all groups. Deconvoluted clonotype frequencies were scaled by the per-sample coverage factor and, where two bulk TCR-seq technical replicates were available, averaged across the two replicates to obtain a single frequency vector per fraction-timepoint combination. For tp3 bulk TCR-seq samples, one replicate had suspected mislabeling of *α* and *β* libraries and removed in the analysis.

Sequencing depth was expressed as the mean number of sequencing reads attributed to each PBMC input, computed as

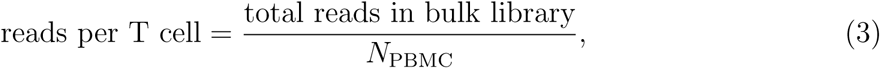

where *N*_PBMC_ = 1.5 *×* 10^6^ input PBMCs. Deconvolution accuracy was assessed against the single-cell TCR-seq ground truth from the matching timepoint (tp1 or tp3 10x data).

## Acknowledgement

This work was supported by the Chinese Academy of Medical Sciences (CAMS) Innovation Fund for Medical Science (CIFMS), China (grant number: 2024-I2M-2-001-1). Y.Z. is supported by China Scholarship Council. We thank Ricardo A. Fernandes from COI and Rachael Bashford-Rogers from CHG for suggestions.

## Author contributions

This project is supervised by T.D., J.C.K., E.A., and Y.P.; Y.Z and E.A. designed the experiments; Y.Z. and J.Z. designed the algorithm; Y.Z. performed the experiments and analyzed the data; M.P.P and E.A. collected and curated the breast cancer data; G.A, F.I, Y.W. collected and processed the melanoma sample; E.A. curated melanoma data; Y.Z., E.A., M.P.P., and Y.P. wrote the manuscript; All authors have read and approved the final manuscript.

## Data availability

ClonoDeconv package is publicly available at https://github.com/YuzeGO/ClonoDeconv. All additional intermediate files are available from the corresponding author upon reasonable request.

## Conflict of interests

The authors declare no conflict of interests.

## Notes

### Competing Interest Statement

The authors have declared no competing interest.

