## Supplemental material for "ClonoDeconv: reference-based *α − β* paired recurrent clonotype proportion estimation from bulk-TCR sequencing"

---

**S2 Supplementary Figures** **6**

### S1 Supplementary Methods

#### S1.1 scTCR data preprocessing

**10x Genomics.** Raw Cell Ranger VDJ files were filtered for `productive`, `full_length`, `is_cell`, and `high_confidence` contigs. Chains were separated by type (TRA, TRB). The within-cell clonotype fraction for chain $j$ in cell $i$ was:

$$f_{i,j} = \frac{c_{i,j}}{\sum_{j'} c_{i,j'}}, \tag{S1}$$

where $c_{i,j}$ denotes the UMI count of chain $j$ in cell $i$ and the sum runs over all chains of the same type. Chains with $f_{i,j} < 0.2$ were discarded to suppress false-positive low-abundance chains.

**SMART-seq.** Read counts from MiXCR output were normalised within each sequencing plate column (batch) $b$:

$$\tilde{c}_{i,j}^{(b)} = \frac{c_{i,j}^{(b)}}{\sum\limits_{i',j' \in b} c_{i',j'}^{(b)}} \times \left|\mathcal{B}_b\right|, \tag{S2}$$

where $\mathcal{B}_b$ is the set of chain records in batch $b$. This normalisation places counts from all batches on the same scale while preserving relative within-batch abundances.

**Clonotype definition.** A single-chain was defined as the string V-gene_CDR3aa. A paired clonotype was defined as the concatenation of the alpha and beta single-chains, separated by "/".

#### S1.2 scTCR reference construction

Let $\mathcal{A}$ and $\mathcal{B}$ denote the sets of unique alpha and beta chain identifiers, respectively, and let $\mathcal{K}$ denote the set of unique paired clonotypes present in the single-cell dataset. The reference matrix $\mathbf{R} \in \mathbb{R}^{(|\mathcal{A}|+|\mathcal{B}|)\times|\mathcal{K}|}$ is constructed so that entry $R_{j,k}$ represents the expected expression level of chain $j$ in cells bearing clonotype $k$.

For cells expressing a single alpha chain paired with one beta chain, the dual-alpha weight is $w_{i,k} = 1$. For cells expressing $d > 1$ alpha chains with a single beta chain, each alpha–beta pairing $(a_\ell, \beta)$ receives weight

$$w_{i,k_\ell} = \frac{c_{i,a_\ell}}{\sum\limits_{\ell'=1}^{d} c_{i,a_{\ell'}}}, \tag{S3}$$

reflecting the fractional UMI (10x) or read (SMART-seq) contribution of each alpha chain. The weights for a given cell sum to one across all pairing records.

The reference entry for chain $j$ and clonotype $k$ is:

$$R_{j,k} = \frac{\sum_{i \in \mathcal{N}_k} w_{i,k}\, c_{i,j}}{\sum_{i \in \mathcal{N}_k} w_{i,k}}, \tag{S4}$$

where $\mathcal{N}_k = \{i : \text{cell } i \text{ bears clonotype } k\}$ is the set of cells expressing clonotype $k$, $c_{i,j}$ is the (normalised) count of chain $j$ in cell $i$, and $w_{i,k}$ is the dual-alpha weight. For UMI-based (10x) data, $c_{i,j}$ are raw UMI counts; for SMART-seq data, $c_{i,j}$ are batch-normalised read counts.

#### S1.3 Finding clonotype clusters and reconstructing the grouped reference

**Graph-based clustering (recommended for large datasets).** Formally, a graph $G = (V, E)$ is constructed where $V$ indexes the rows of $\mathbf{R}$ (unique chains), and an edge $(u, v)$ is added for every clonotype $k$ in which both chain $u$ (alpha) and chain $v$ (beta) are non-zero — i.e., the edge encodes the pairing. Connected components of $G$ define clonotype clusters: clonotypes within the same component share at least one chain with another member and therefore cannot be independently resolved by the bulk signal alone without additional constraints.

**Distance-based clustering (for smaller datasets).** For datasets with fewer clonotypes (generally less than 200 clones, as too many clones will cause problems in dentrogram cutting), hierarchical clustering could be performed on a binary clonotype–chain incidence matrix using Jaccard distance and average linkage. A distance threshold is selected from the resulting dendrogram to define flat clusters.

**Grouped reference reconstruction.** Given a cluster assignment mapping $\phi : k \mapsto g$, the grouped reference matrix $\mathbf{R}_g \in \mathbb{R}^{(|\mathcal{A}|+|\mathcal{B}|)\times|\mathcal{G}|}$ is constructed by aggregating per-clonotype contributions within each cluster $g$:

$$R_{g,j,g'} = \frac{1}{N_{g'}} \sum_{k:\phi(k)=g'} \sum_{i \in \mathcal{N}_k} c_{i,j}, \quad j \in \mathcal{A} \cup \mathcal{B}, \tag{S5}$$

where $N_{g'} = |\bigcup_{k:\phi(k)=g'} \mathcal{N}_k|$ is the number of cells in cluster $g'$. Cluster *markers* $\mathcal{M}_g$ are defined as the chains with non-zero entries in $\mathbf{R}_g[\cdot, g]$ for clusters containing more than one clonotype; these are the only chains that carry discriminating information for the refined second stage.

#### S1.4 Recursive non-negative least squares (NNLS) estimation of clonotype frequencies

**Stage 1 – grouped NNLS.** Let $\mathbf{b} \in \mathbb{R}^M$ be the bulk count vector ($M = |\mathcal{A}| + |\mathcal{B}|$). Clonotype-group proportions are estimated by solving:

$$\hat{\mathbf{x}}_g = \arg\min_{\mathbf{x}_g \geq \mathbf{0}} \big\|\mathbf{R}_g\, \mathbf{x}_g - \mathbf{b}\big\|_2^2. \tag{S6}$$

The problem is solved using Fast Iterative Shrinkage-Thresholding (FISTA-projected gradient descent) implemented in PyTorch with optional GPU acceleration.

**Stage 2 – refined clonotype-level NNLS.** For singleton clusters ($|\mathcal{C}_g| = 1$), the refined estimate equals the grouped estimate directly. For multi-member clusters ($|\mathcal{C}_g| > 1$), the optimisation is restricted to the marker chains $\mathcal{M}_g$ and the clonotypes in $\mathcal{C}_g$, and the optimisation target is constructed to isolate the signal attributable to that cluster.

Define the cluster's estimated relative contribution:

$$p_g = \frac{\hat{x}_g}{\sum_{g'} \hat{x}_{g'} + \epsilon}, \tag{S7}$$

where $\epsilon = 10^{-10}$ avoids division by zero. For each marker chain $j \in \mathcal{M}_g$, the refined target is:

$$t_{c,j} = R_{j,g}\,\hat{x}_g + \left(b_j - \mathbf{r}_j^\top \hat{\mathbf{x}}_g\right) p_g, \tag{S8}$$

where $\mathbf{r}_j$ is the $j$-th row of the grouped reference $\mathbf{R}_g$. The first term scales the expected expression from the cluster alone; the second redistributes the residual bulk signal at chain $j$ proportionally to the cluster's estimated size. This construction ensures that only the fraction of the bulk signal plausibly attributable to cluster $g$ enters the within-cluster NNLS, and that cross-cluster bleed-in is reduced.

The within-cluster NNLS is then:

$$\hat{\mathbf{x}}_{\mathcal{C}_g} = \arg\min_{\mathbf{x} \geq \mathbf{0}} \left\| \mathbf{R}\left[\mathcal{M}_g, \mathcal{C}_g\right] \mathbf{x} - \mathbf{t}_c \right\|_2^2, \tag{S9}$$

where $\mathbf{R}[\mathcal{M}_g, \mathcal{C}_g]$ is the sub-matrix of the original reference restricted to the marker chains and the clonotypes of cluster $g$. Restricting the problem to marker chains exploits the fact that only chains present in the cluster reference encode discriminating information; chains shared by all members of a cluster or not expressed by any member contribute no resolution and are excluded.

#### S1.5 Bulk-TCR dataset handling

**Coverage factor.** Let $\mathcal{S}$ denote the set of chain identifiers present in the reference $\mathbf{R}$. The coverage factor for a bulk sample $\mathbf{b}$ is:

$$f_{\text{cov}} = \frac{\sum\limits_{j \in \mathcal{S}} b_j}{\sum\limits_{j} b_j}, \tag{S10}$$

representing the fraction of total bulk signal that can be attributed to chains present in the reference. Clonotypes absent from the scTCR-seq reference — either because they were not captured in single-cell sequencing or because they are private to the bulk sample — contribute to the denominator but not the numerator, giving $f_{\text{cov}} < 1$ in typical experiments.

**Bulk trimming.** Before deconvolution, the bulk vector is trimmed to the reference index: chains in the bulk that are not present in the reference are removed, and chains present in the reference but absent from the bulk are set to zero. The coverage factor is stored and applied after deconvolution.

**Proportion scaling.** After the two-stage NNLS, the raw estimates $\hat{\mathbf{x}}$ are scaled so that their sum equals the coverage factor:

$$\hat{p}_k = \hat{x}_k \cdot \frac{f_{\text{cov}}}{\sum_{k'} \hat{x}_{k'} + \epsilon}. \tag{S11}$$

This anchors the total estimated repertoire fraction to the portion of the bulk that can be explained by the reference, preserving the interpretation that $1 - f_{\text{cov}}$ corresponds to the contribution of clones not present in the reference. Numerical accuracy is improved because the sum constraint prevents the solver from absorbing background noise into arbitrarily large aggregate proportions.

### S2 Supplementary Figures

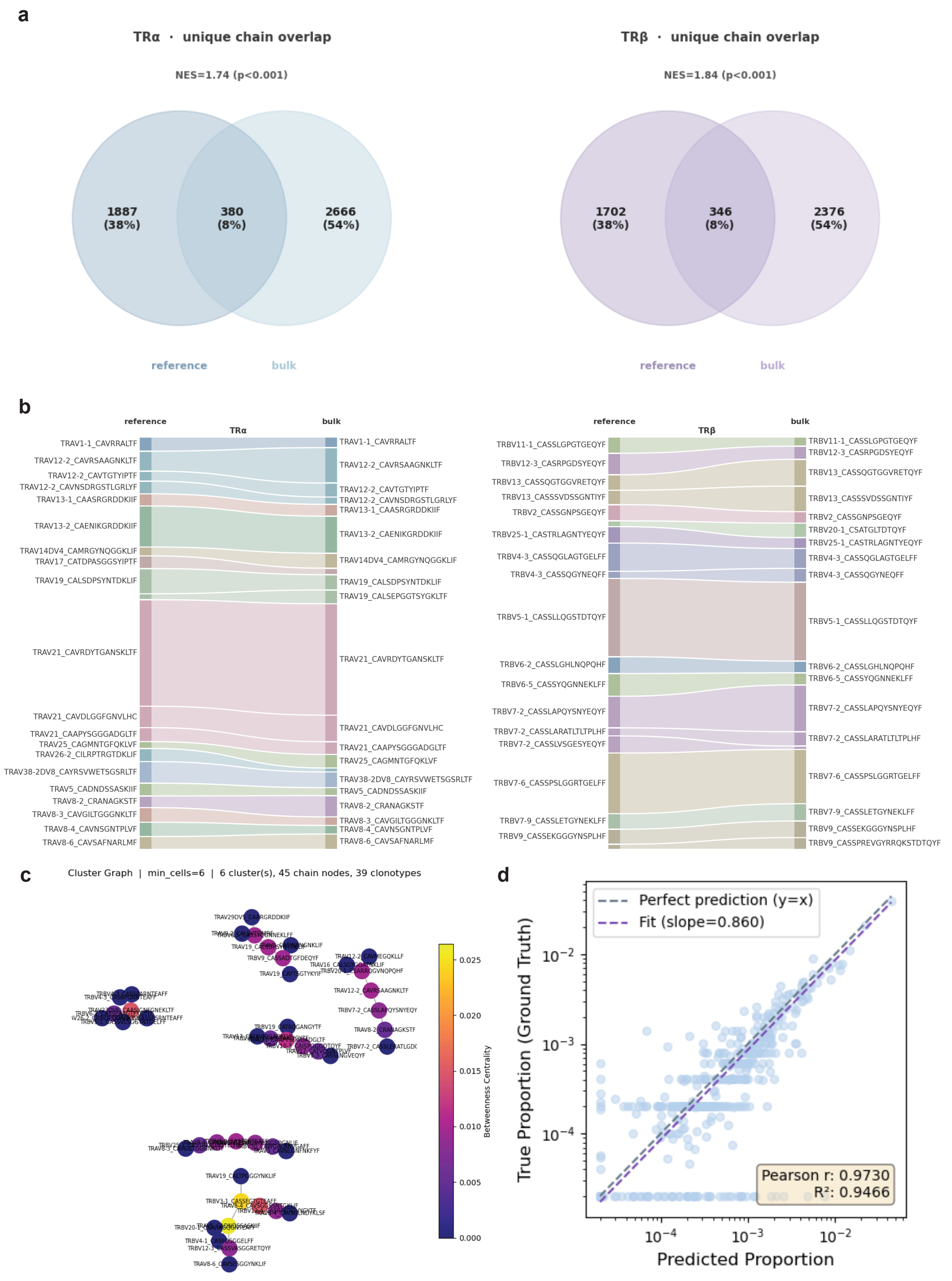

**Supplementary Figure 1. Alignment of 10x scTCR pseudobulk and paired reference a**, Venn plot showing the number of overlaping unique $\alpha$ (left) and $\beta$ (right) chains between the pseudobulk and the reference, NES and p values were calculated with GSEApy (Fang et al., *Bioinformatics*, 2023) to evaluate the enrichment of reference chains in pseudobulk. **b**, Sankey plot showing the union of top 15 single chains in reference and pseudobulk, left=$\alpha$ and right=$\beta$; **c**, Representative 6 largest clonotype groups after graph-based clonotype clustering. Nodes represent $\alpha$ or $\beta$ single chains, edges linking two nodes represent $\alpha\beta$ paired clonotypes, colour denotes betweenness centrality; **d**, Scatter plot comparing deconvoluted clonotype frequencies of individual clonotypes vs. ground truth frequencies in OX1b, blue line shows $y = x$, purple dotted line is regression line, $R^2$ is the coefficient of determination.

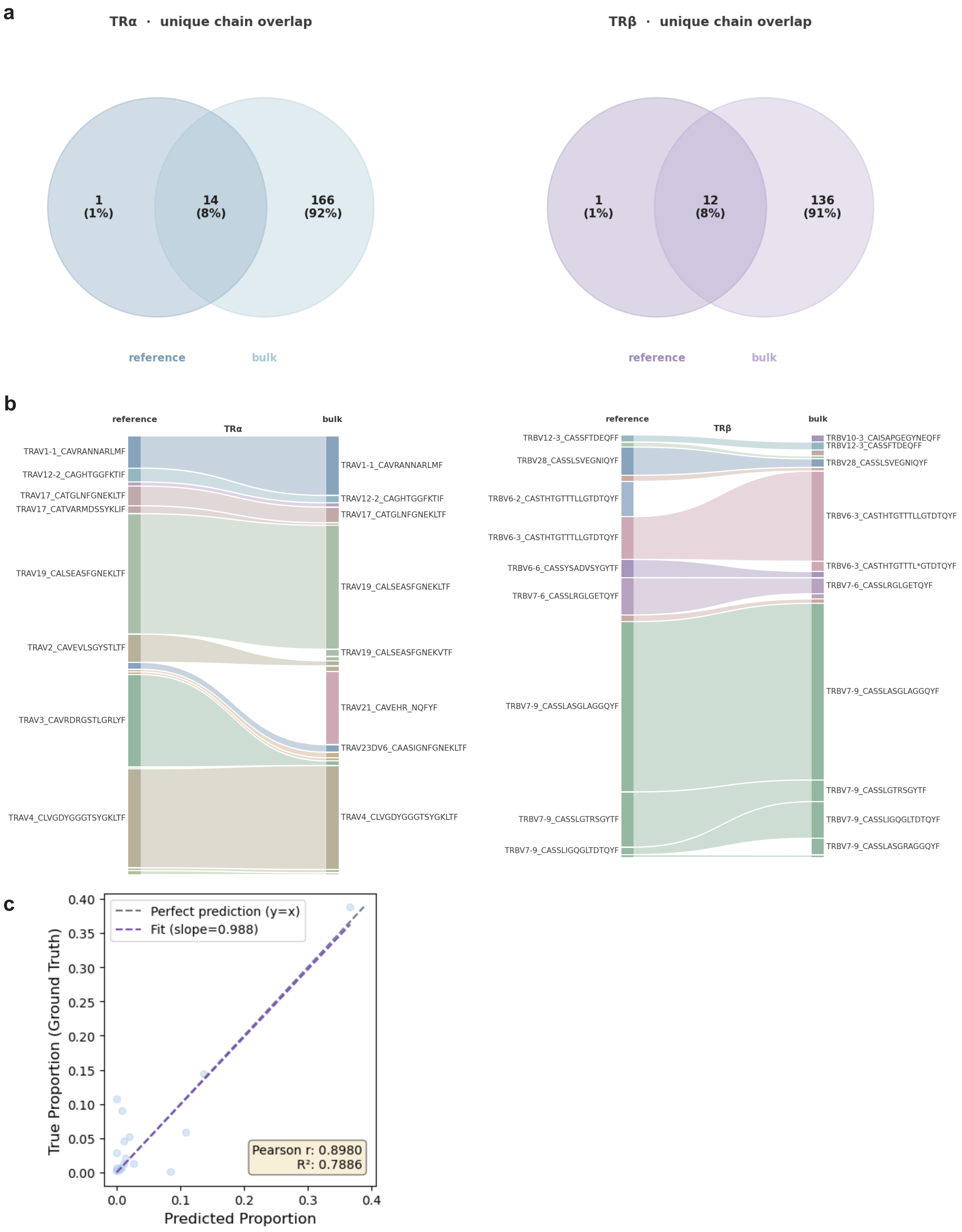

**Supplementary Figure 2. Alignment of SMART-seq scTCR and paired bulk-TCR profile of tumour-specific T cells. a**, Venn plot showing the number of overlaping unique $\alpha$ (left) and $\beta$ (right) chains between the bulk and the reference; **b**, Sankey plot showing the union of top 15 single chains in reference and bulk, left=$\alpha$ and right=$\beta$; **c**, Scatter plot comparing deconvoluted clonotype frequencies of individual clonotypes vs. ground truth frequencies, blue line shows $y = x$, purple dotted line is regression line, $R^2$ is the coefficient of determination.

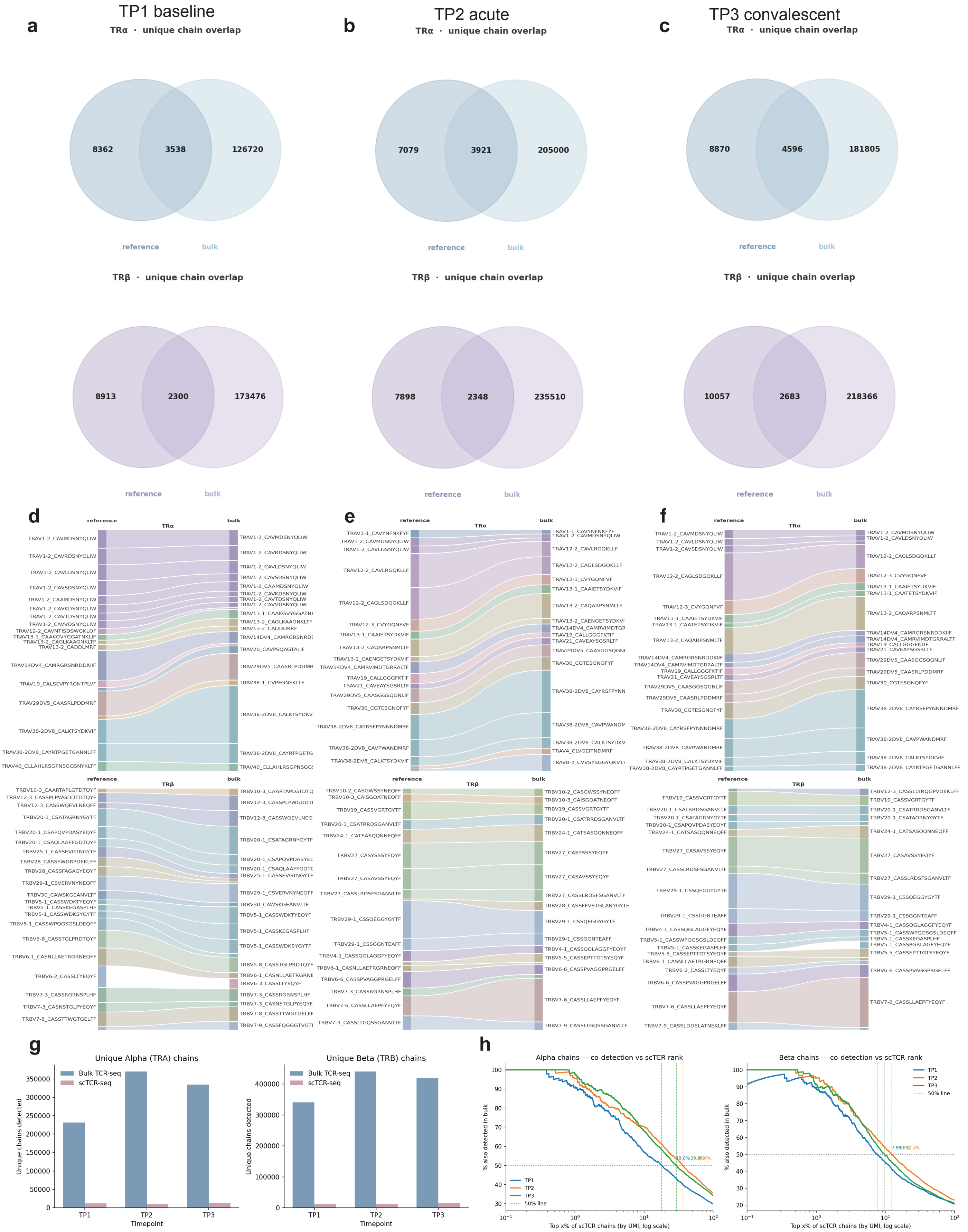
a
TP1 baseline
TRα · unique chain overlap
8362
3538
126720
reference
bulk
TRβ · unique chain overlap
8913
2300
173476
reference
bulk
b
TP2 acute
TRα · unique chain overlap
7079
3921
205000
reference
bulk
TRβ · unique chain overlap
7898
2348
235510
reference
bulk
c
TP3 convalescent
TRα · unique chain overlap
8870
4596
181805
reference
bulk
TRβ · unique chain overlap
10057
2683
218366
reference
bulk
d
e
f
reference
TRα
bulk
TRβ
g
Unique Alpha (TRA) chains
Unique Beta (TRB) chains
Bulk TCR-seq
scTCR-seq
Unique chains detected
Timepoint
TP1
TP2
TP3
h
Alpha chains — co-detection vs scTCR rank
Beta chains — co-detection vs scTCR rank
% also detected in bulk
Top x% of scTCR chains (by UMI, log scale)
TP1
TP2
TP3
50% line

**Supplementary Figure 3. Alignment of 10x scTCR and paired bulk-TCR profile of longitudinal samples.** **a**, **b**, **c**, Venn plot showing the number of overlaping unique $\alpha$ (top) and $\beta$ (bottom) chains between the bulk and the reference at tp1/baseline (**a**), tp2/acute (**b**), and tp3/convalescent (**c**); **d**, **e**, **f**, Sankey plot showing the union of top 15 single chains in reference and bulk, top=$\alpha$ and bottom=$\beta$ at tp1/baseline (**d**), tp2/acute (**e**), and tp3/convalescent (**f**); **g**, Number of detected unique $\alpha$ (left) and $\beta$ (right) chains at each timepoint from bulk-TCR (blue) and scTCR (purple) sequencing; **h**, Percentage of unique single chains in top x% of scTCR that are co-detected in paired bulk, x-axis denotes top x% of all $\alpha$ (left) and $\beta$ (right) chains in scTCR, y-axis denotes percentage of overlapped chains with bulk-TCR in that top x% of chains. Horizontal dotted line represents 50% of overlap, vertical dotted lines and numbers are the corresponding percentage in scTCR that could achieve such 50% of overlap, blue=tp1, orange=tp2, green=tp3.

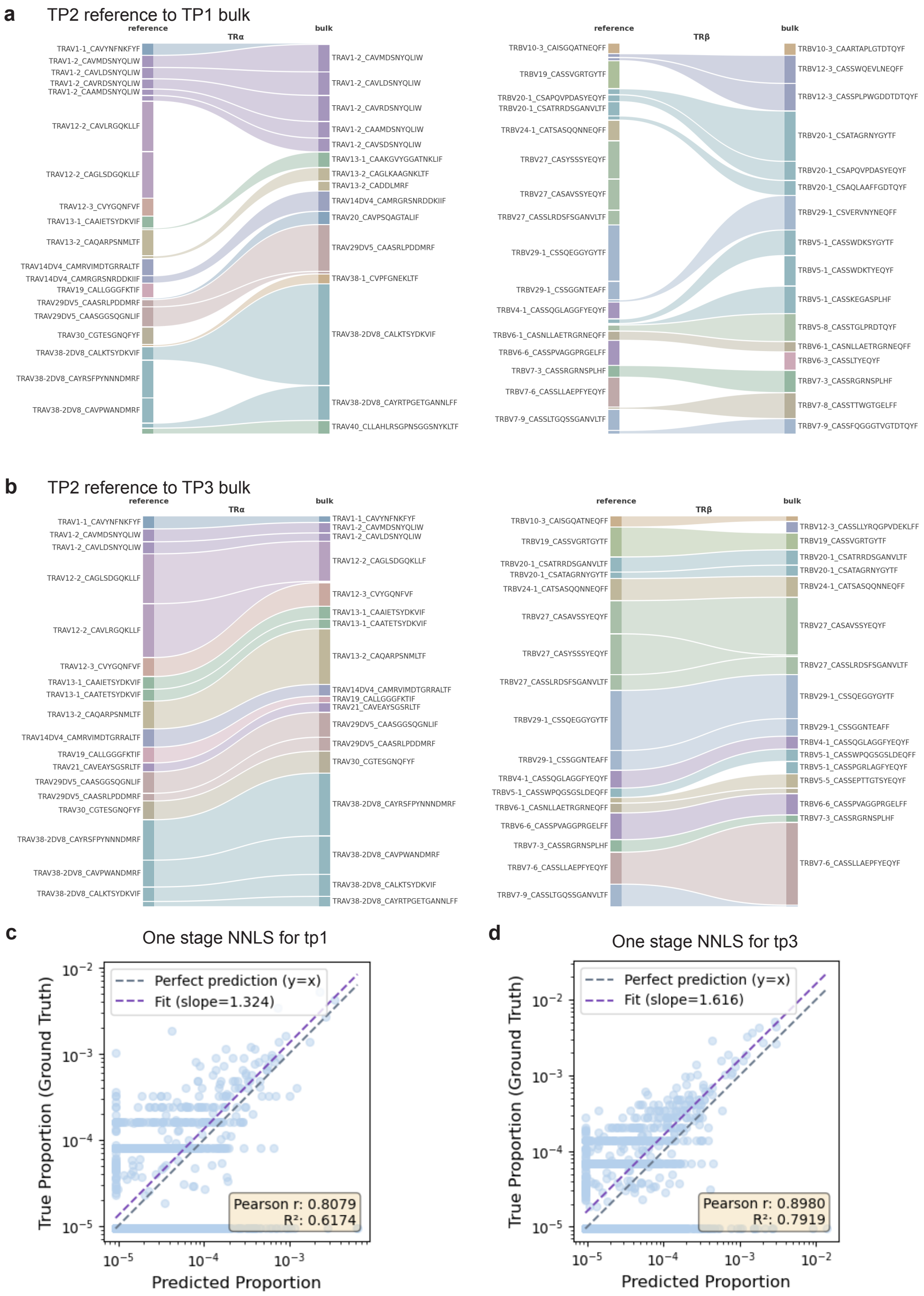

**Supplementary Figure 4. Cross-timepoint alignment between reference and bulk and simple NNLS deconvolution. a**, **b**, Sankey plot showing the union of top 15 single chains in tp2 reference and tp1 bulk (**a**) or tp3 bulk (**b**), left=$\alpha$ and right=$\beta$; **c**, **d**, Scatter plot comparing deconvoluted clonotype frequencies of individual clonotypes vs. ground truth frequencies at tp1 (**c**) and tp3 (**d**), blue line shows $y = x$, purple dotted line is regression line, $R^2$ is the coefficient of determination.

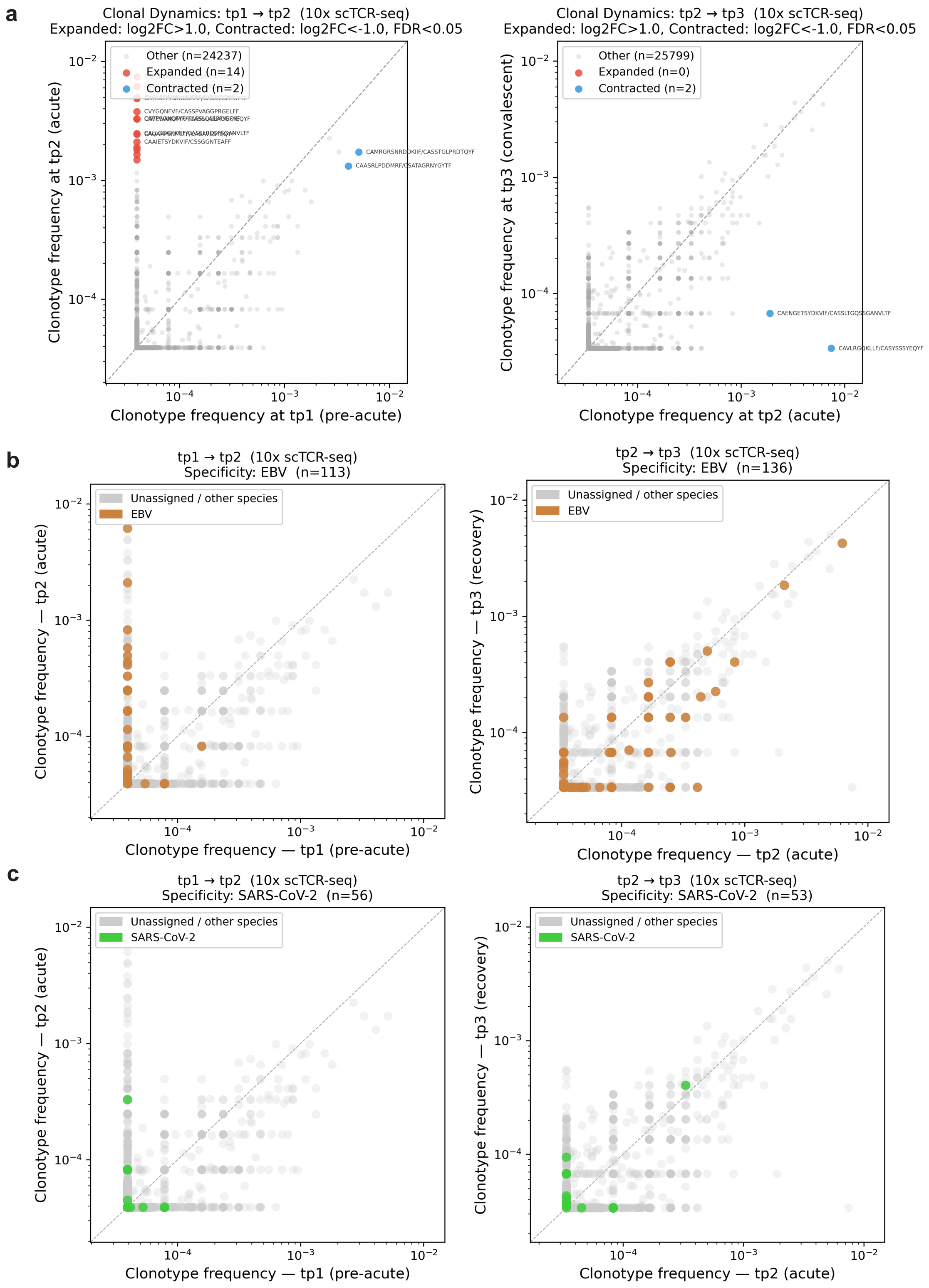

**Supplementary Figure 5. Tracking $\alpha\beta$ paired clonal dynamics in longitudinal bulk-TCR samples with 10x scTCR at all timepoints.** **a**, Identification of significantly expanded (red) and contracted (blue) clones between pairwise timepoints (left, tp1 to tp2; right, tp2 to tp3). Significance given by log2 fold change >1 or < -1, and FDR < 0.05 (Fisher exact test with BH multiple comparison correction). All other non-significant clonotypes are in grey; **b**, Clonal dynamics of EBV-specific clones (yellow) between pairwise timepoints (left tp1-tp2, right tp2-tp3). Frequencies at all timepoints are given by 10x scTCR; **c**, Clonal dynamics of SARS-CoV-2 specific clones (green) on pairwise timepoints (left tp1-tp2, right tp2-tp3). Frequencies at all timepoints are given by 10x scTCR.
